# Post-flowering photoperiod sensitivity of soybean in pod-setting responses

**DOI:** 10.1101/2024.06.03.597100

**Authors:** Zhihui Sun, LiMei Yuan, Yulin Wang, Ran Fang, Xiaoya Lin, Haiyang Li, Liyu Chen, Yichun Wu, Xin Huang, Fanjiang Kong, Baohui Liu, Sijia Lu, Lingping Kong

## Abstract

The development of soybean (*Glycine max*) is regulated by photoperiod, with genes related to photoperiod sensitivity primarily focused on flowering time. However, their roles in post-flowering reproductive development and the mechanisms by which photoperiod affects them are not yet determined. In this study, we found that pod formation is sensitive to photoperiod. Long-day (LD) conditions tend to extend the time from flowering to pod formation (R1 to R3 stage), and the first wave of flowers tends to fall off. Additionally, photoperiod affects pistil morphology; under short-day (SD) conditions, the stigma has a curved hook-like structure that facilitates better interaction with the filaments when pollen is released, ultimately influencing the timing of pod formation. Photoperiod-insensitive mutants, lacking *E1* family and *Evening Complex* genes, showed no difference in pod formation time under LD or SD conditions. Hormone content analysis and transcriptome data analysis indicated that various hormones, ROS signals, and the application of sucrose solution *in vitro* might influence floral organ abscission.

**Highlight:** Photoperiod sensitivity after flowering affects the pod-setting time in soybean.

## Introduction

Photoperiod is a rhythmic change in the amount of light received by an organism. Plants sense photoperiod, which enables them to adjust their flowering time according to seasonal changes in light to adapt to growing conditions at different latitudes (Garner and Allard, 1920; Hayama *et al*., 2003; Silva *et al*., 2020; Jung *et al*., 2020; Bu *et al*., 2021). In addition to regulating flowering time, photoperiod also affects physiological processes such as photosynthesis, growth rhythm, and nutrient metabolism of plants. Plants adjust the intensity and time of photosynthesis by sensing the photoperiod to maximize the use of light energy for nutrient synthesis and growth development. Photoperiod is also closely related to processes such as the distribution of photosynthetic products, carbon metabolism, and the synthesis of phytohormones, directly affecting the growth rate and morphological structure of plants.

Soybean is a typical short-day (SD) crop, which is very sensitive to changes in photoperiod. Usually, one variety or germplasm resource is suitable for planting in a particular narrow latitude range because modern cultivated soybean varieties require such specific photoperiods (Watanabe *et al*., 2012; Lu *et al*., 2017). The wide genomic adaptability of soybean is mainly achieved through changes in the multiple genes or quantitative trait loci that control the flowering and reproductive period. A growing number of photoperiod-responsive gene loci have been identified and analyzed at the molecular level, including the *E* series (*E1*–*E4*, *E9*) and *Time of flowering* (*Tof5*), *Tof11*, *Tof12*, *Tof16*, *LUX ARRHYTHMO* (*Lux*), and *J* (Liu *et al*., 2008; Watanabe *et al*., 2009; Watanabe *et al*., 2011; Xia *et al*., 2012; Kong *et al*., 2010; Kong *et al*., 2014; Zhao *et al*., 2016; Lu *et al*., 2017; Lu *et al*., 2020; Bu *et al*., 2021; Dong *et al*., 2021; Dong *et al*., 2022). The flowering time loci *E1*, *E2*, *E3, E4*, *Tof11,* and *Tof12* play a role in regulating long-day (LD) insensitivity (where mutants of these genes tend to flower earlier even under non-inductive LD conditions such as high latitudes.) (Lu *et al*., 2020; Xu *et al*., 2013; Lu *et al*., 2020). Over 80% of low-latitude soybean varieties harbor different mutant alleles in the *J* and *Tof16* genes, suggesting that *Tof16* and *J* play a significant role in soybean adaptation to SD photoperiods (Mutants of these genes tend to have long juvenile and flower late under induced SD (low latitude) condition) (Dong *et al*., 2021).

The photoperiodic response of soybeans not only operates during the pre-flowering growth stages but also plays a crucial role in post-flowering vegetative and reproductive growth processes. (Kantolic *et al*., 2007; Jiang *et al*., 2010; Xu *et al*., 2013; Kim *et al*., 2020). During the post-flowering stages, plants remain sensitive to photoperiod, and this sensitivity is also regulated by maturity genes. (Summerfield *et al*., 1998; Ellis *et al*., 2000). The interaction between genes and the environment that control the reproductive period directly affects various phenotypic characteristics in the post-flowering stages, such as pod-setting, pod development, terminal vegetative growth, and reproductive growth (Curtis *et al*., 2000; Kantolic and Slafer 2001, 2005, 2007; Cooper *et al*., 2003; Xu *et al*., 2013; Nico *et al*., 2016). The extended duration of R3-R6 under longer photoperiods tend to increase pod and seed number (Kantolic and Slafer 2007). Post-flowering photoperiod extension delayed individual fruit development in soybean from R1 stage to seed filling stage (Nico *et al*., 2016). However, while we have observed the influence of photoperiod on the flowering to pod setting process, the molecular mechanisms involved remain unclear. Apart from maturity genes, genes potentially involved in regulating the flowering to pod setting process may include those related to light signal transduction, plant hormone regulation, carbon metabolism, and nutrient transport. These genes interact through complex signaling networks, regulating soybean growth, development, and yield formation during the post-flowering stages. In-depth studies of these genes can help us comprehensively understand the growth regulatory mechanisms of soybeans, providing scientific basis for improving soybean yield and quality.

Long days lengthened the flowering period and thereby increased the number of opened flowers on lateral racemes. During the post-flowering phase, seed filling effectiveness was delayed on primary racemes (dominant positions), enhancing the pod number on lateral racemes (usually dominated positions) at some main stem nodes in long day conditions (Nico *et al*., 2016). This phenomenon is often observed under artificial light conditions in greenhouses or growth chambers: under long-day conditions (e.g. 16 hours light/8 hours dark), the first flowers to bloom of most soybean varieties gradually fall off instead of developing into pods. In contrast, under artificial short-day conditions (e.g. 12 hours light/12 hours dark), flowers begin to produce pods more quickly. Prolonged daylight hours also delay the time for soybean flowers to develop into pods, extending the pod initiation period without altering the rate of pod elongation (Nico *et al*., 2016). This indicates the influence of photoperiod on the pod development process while also suggesting the potential involvement of other factors affecting pod development and maturation. There is a complex relationship between pod abscission and photoperiodic responses. Environmental stresses such as low light radiation conditions are important factors that may induce flower buds abscission (Ren *et al*., 2022). Studies in different species have shown that flower/fruit abortion is determined by the availability of assimilates (Marcelis *et al*., 2004; Ali *et al*., 2022; Ren *et al*., 2022). When seeds enter the linear phase of growth and accumulate assimilates at their maximum rate, they become a relatively large reproductive sink that may limit upcoming flowering, resulting in flower abortion to allow the older organs to finish their development (Turc *et al*., 2018). Sugar signaling plays a potential central role in regulating lotus (*Nelumbo nucifera*) flower bud abortion; for example, the overexpression of *Trehalose-6-P Synthase 1* (*TPS1*) in lotus significantly decreased the flower bud abortion rates in both normal-light and low-light environments (Ren *et al*., 2022). This illustrates the importance of sugar signals in regulating post-anthesis development, possibly affecting soybean pod development and maturation by regulating the distribution and utilization of assimilates. It is proposed that flower abortion could be mediated by hormonal induction, potentially by the candidate hormone indole-3-acetic acid (IAA) (Huff and Dybing, 1980). Abscisic acid could also be involved because it has an inhibitory role on flowering (Bernier *et al*., 1993).

Flower and pod abscission are important factors affecting soybean crop yields. Therefore, analyzing the physiological mechanisms of photoperiodic regulation on flowering and subsequent pod development is of significant importance for promoting crop breeding and genetic improvement. In this study, we observed that the time from flowering to pod formation on the whole soybean plant was longer under LD conditions than under SD conditions. Such differences under different photoperiods were not observed in photoperiod-insensitive soybean genotypes, indicating that the period between flowering and pod setting is sensitive to day length. Furthermore, we found the pod-setting signal is mainly induced and transmitted by leaves. We therefore showed that photoperiod affects the various stages of soybean growth and development. Further research into the molecular mechanisms regulating the time between flowering and pod setting will be helpful for improving soybean yields through the reduction of flower and pod abortion.

## Materials and methods

### Plant materials, growth conditions, and phenotyping

The soybean cultivars Williams 82 (W82; *e1-as/E2/E3/E4*) (Kong *et al*., 2018) and Harosoy (*e1-as/e2/E3/E4*) (Xia *et al*., 2012) were used in this study. W82 is more sensitive to long photoperiods than Harosoy. Using W82 as the wild type, homozygous transgenic *lux* double mutants (*lux-2m*; *lux1 lux2-2* as published (Bu *et al*., 2021)), and *e1* triple mutants (*e1-3m; e1/e1la/e1lb* mutant type as described (Lin *et al.,* 2022)), and the wild-type plants were used for the experiments. Plants were grown under artificial SD (12 h light/12 h dark), artificial LD (16 h light/8 h dark) and ultra-long day (20 h light/4 h dark) conditions in a greenhouse or a growth chamber, with a light intensity of 240 µmol m^-2^ s^-1^ and a temperature of 25°C. According to the description of the developmental stages of soybean (Fehr *et al*., 1971), the reproductive stages R1 and R2 are based on flowering, R3 and R4 on pod development, R5 and R6 on seed development, and R7 and R8 on maturation. Flowering time was recorded at the R1 stage as the number of days from seedling emergence to the first open flower at any node on the main stem. The pod-setting time was recorded when any node at the four upmost produced a pod with a length of 0.5 cm. At least five plants were detected for each line.

### Transfer between different photoperiod conditions

W82 plants were grown under LD (16 h light/8 h dark) conditions until R1 in the green house, after which half were transferred into SD (12 h light/12 h dark) conditions (named LD_SD group) with the others remaining in the LD (16 h light/8 h dark) treatment (named continuous LD group or LD_LD group). Pod setting time were then measured after the transferred treatments. The duration of these treatment was 60 days.

### Branch-specific photoperiod treatments

W82 plants were grown under LD (16 h light/8 h dark) conditions until the fifth day after emergence in the growth chamber. To ensure branching in each plant, the shoot apical meristems (SAMs) were cut to remove the apical dominance and promote the development of lateral branches. All plants were grown in LD conditions until reaching the R1 stage after which the branches were subjected to treatments of different photoperiodic combinations. In one set of experiments, the light phase of one branch was shortened to 12 hours using black bags to exclude light. The bags were removed each day and then replaced at Zeitgeber time 12 (ZT 12) each day. When the light was on (at ZT 0) they were removed. To remove any phenotypic differences caused by this bagging, another branch was covered with transparent plastic bags as the LD control. In another set of experiments, both branches were covered with transparent plastic bags and subjected to LD conditions. To further demonstrate the role of leaves in perceiving the photoperiod and controlling the pod initiation time, all leaves of branches under different photoperiod conditions were removed, with another branch retaining its leaves as the control.

### Pollen germination analysis

The pollen germination experiments were based on in vitro and in vivo pollen germination. In brief, for in vitro pollen germination, mature pollen grains of W82 under LD and SD conditions were dispersed on pollen germination medium containing 10% sucrose, 0.01% boric acid, 5 mM CaCl_2_, 5 mM KCl, 1 mM MgSO_4_, pH 7.5 and 1.5% agar (Boavida and McCormick 2007). Germination mediums were then incubated at 25°C temperature for 7 hours. Pollen germination was observed under microscope (Zeiss Axio Imager A2). Pollen tube length was measured by ImageJ software (Version 1.8.0). For *in vivo* germination experiments, pollen grains were applied on stigmata of W82 *under LD and SD conditions.* After 20 hours, the hand-pollinated pistils were fixed in a solution of 45%:6%:5% acetic acid/ethanol/formaldehyde for 2 hours, washed with 70% ethanol, 50% ethanol, 30% ethanol and ddH_2_O for 10 minutes each, and then treated with 8 M NaOH overnight. Samples were washed three times with ddH_2_O and stained with aniline blue solution (0.1% aniline blue, 108 mM K_3_PO_4_) for more than 2 hours (Mori *et al*., 2006). Stained samples were observed under a fluorescence microscope (Zeiss Axio Imager A2).

### Transcriptome analysis

Flower buds samples before flowering were collected at Zeitgeber time 4 at R1 stage under LD (16 h light/8 h dark) and SD (12 h light/12 h dark) conditions of W82, with each sample collected from 5 individual plants. Analysis was conducted on three biological replicate samples. Pistils were detached from the pod. Experimental methods for total RNA extraction, Illumina sequencing, and RNA differential expression analysis were performed following procedures described in previous publications (Bu *et al*., 2021). Genes/transcripts with false discovery rate (FDR) values below 0.05 and absolute fold change ≥ 2 were considered as differentially expressed genes/transcripts. Soybean reference genome used in this study including https://www.ncbi.nlm.nih.gov/datasets/genome/GCF_000004515.6/ and https://phytozome-next.jgi.doe.gov/info/Gmax_W82_a4_v1.

### Quantitative reverse-transcriptase (RT)-PCR

Total RNA was extracted from pistils of flower buds before opening, at R1 stage and 1 day, 5days and 10 days post R1 stage in LD_LD and LD_SD groups using TRIzol regent (Invitrogen). Total RNA was reverse transcribed to cDNA with M-MLV reverse transcriptase kit (Takara). LightCycler 480 SYBR Green I Master (Roche) was used for Quantitative RT-PCR (qRT-PCR) on a Roche LightCycler 480 system (Roche). *Tubulin* was used as an internal control gene. Three biological replications were performed in each test. Primers are listed in Supplementary table 4.

### Phytohormones detection

Phytohormones contents of flower buds of W82 under LD and SD photoperiod conditions were detected by MetWare (http://www.metware.cn/) based on the AB Sciex QTRAP6500 LC-MS/MS platform.

### Sucrose solution spray after R1 stage

W82 plants grown under LD conditions were sprayed with 50mg/ml sucrose solution on their leaves at the R1 stage for 20 days. The blank control group sprayed water without added sucrose, and the pod initiation stage (R3) of the two treatment groups was observed.

### Pathway Enrichment Analysis

Pathway-based analysis helps to further understand genes biological functions. Kyoto Encyclopedia of Genes and Genomes (KEGG) (Kanehisa *et al*., 2000) is the major public pathway-related database (Robinson *et al*., 2010). Pathway enrichment analysis identified significantly enriched metabolic pathways or signal transduction pathways in differently expressed genes (DEGs) comparing with the whole genome background. The calculating formula of *P*-value is:

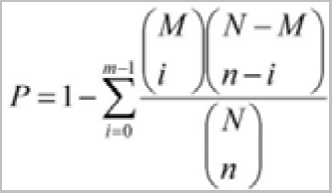

Here N is the number of all genes that with KEGG annotation, n is the number of DEGs in N, M is the number of all genes annotated to specific pathways, and m is number of DEGs in M. The calculated *P*-value was gone through FDR Correction, taking FDR ≤ 0.05 as a threshold. Pathways meeting this condition were defined as significantly enriched pathways in DEGs.

## Results

### Photoperiod affects the initiation of pod-setting after flowering

Under artificial SD (12 h light/12 h dark) and LD (16 h light/8 h dark) conditions, we investigate the flowering time (R1) and the initiation time of podding (R3) of the two cultivars W82 and Harosoy. The time interval between flowering and pod setting initiation (R3-R1) varied among different varieties (Figures 1a-1b). Under LD conditions, successful pod setting typically took approximately 15-30 days after R1 (approximately 15 days for Harosoy and approximately 30 days for W82) (Figures 1a-1b). Comparing the time to pod formation under LD and SD conditions, the trends were similar among different varieties, indicating that pod formation takes significantly longer under LD conditions compared to SD conditions (Fig.1 and Fig.S1). By contrast, under the SD conditions, most of the first-opened flowers successfully initiated pod setting just about three days after R1 (Fig. 1a-1c, Fig.S1c, and Fig. S2a). These results indicate that photoperiod affects the pod-setting time after flowering. Why does soybean require more time to initiate pod setting under LD conditions? We found that under LD conditions, the first-round opened flowers of W82 gradually fell off at most nodes, but later buds continued to be produced; these second-round opened flowers gradually developed into pods. Approximately 16 days after R1, most buds fall off from the nodes on the main stem (Fig. 1b, Fig. S2b). This is one of the reasons for the longer time interval between flowering and pod setting under LD conditions.

**Fig.1 Photoperiod.**
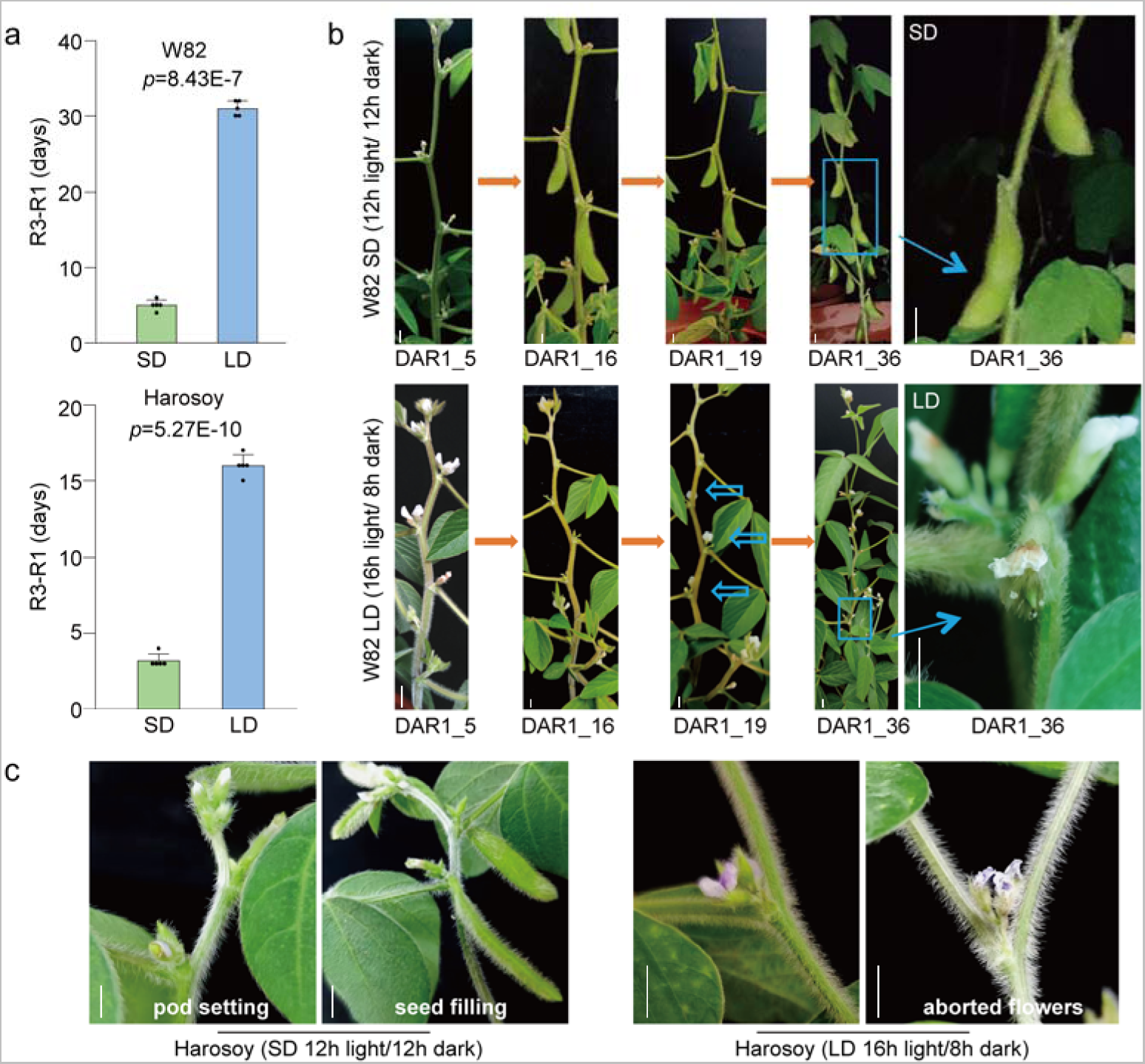
affects R3 stage of different soybean cultivars. (a) The initiation time of pod setting after flowering (duration of R1 to R3 stage) of W82 and Harosoy under short-day (SD; 12 h light/12 h dark) and long-day (LD; 16 h light/8 h dark) photoperiod conditions. (b) Phenotypes of flowers and pods of W82 at 5, 16, 19 and 36 days after flowering under SD and LD condition, respectively. In LD condition, the firstly-opened flowers of W82 gradually fall off instead of developing into pods (DAR1=16). New buds are then produced at each node (DAR1=19), and the second-wave opened flowers (blue arrows) gradually turn into pods. At about 30 DAR1, the pod-setting stage had just begun under long-day conditions, while the pods had reached the seed-filling stage under the short-day conditions (blue dashed box). Under SD, most of the flowers, including many of the first-opened flowers, can successfully initiate pod setting, at about five days after flowering (DAR1=5). (c) Phenotypes of flowers and pod of Harosoy at DAR1=3 and DAR1=8. All data are given as means ± s.e.m. (n = 5 plants). One-tailed, two-sample t-tests were used to generate the *P* values. DAR1, days after R1. The bar in the picture represents 0.5 cm.

### Soybean remains photoperiod-sensitive after flowering

Soybean is known to be sensitive to photoperiod before flowering (Liu *et al*., 2008; Xia *et al*., 2012; Bu *et al*., 2021; Lin *et al*., 2022; Zhao *et al*., 2024); however, the post-flowering sensitivity and mechanisms remain unclear. We grew the soybean cultivar W82 under LD (16 h light/8 h dark) and SD (12 h light/12 h dark) conditions, investigated its phenotypes at R1, R3 and mature stage. W82 displayed different flowering times and plant architectures under different photoperiods. Under SD conditions, plants were smaller with fewer nodes, branches, and pods (Fig. 2a–c). During the period from flowering (R1) to post-flowering (R3), the plants under LD conditions gained about 10 nodes, while those under the SD conditions only gained two nodes during this period (Fig. 2a). These observations indicate that post flowering photoperiod sensitivity not only affects the timing of pod initiation, but also affects plant architecture traits such as node number. Does the significant difference in pod formation rate between LD and SD conditions solely result from differences in plant architectures?

**Fig. 2.**
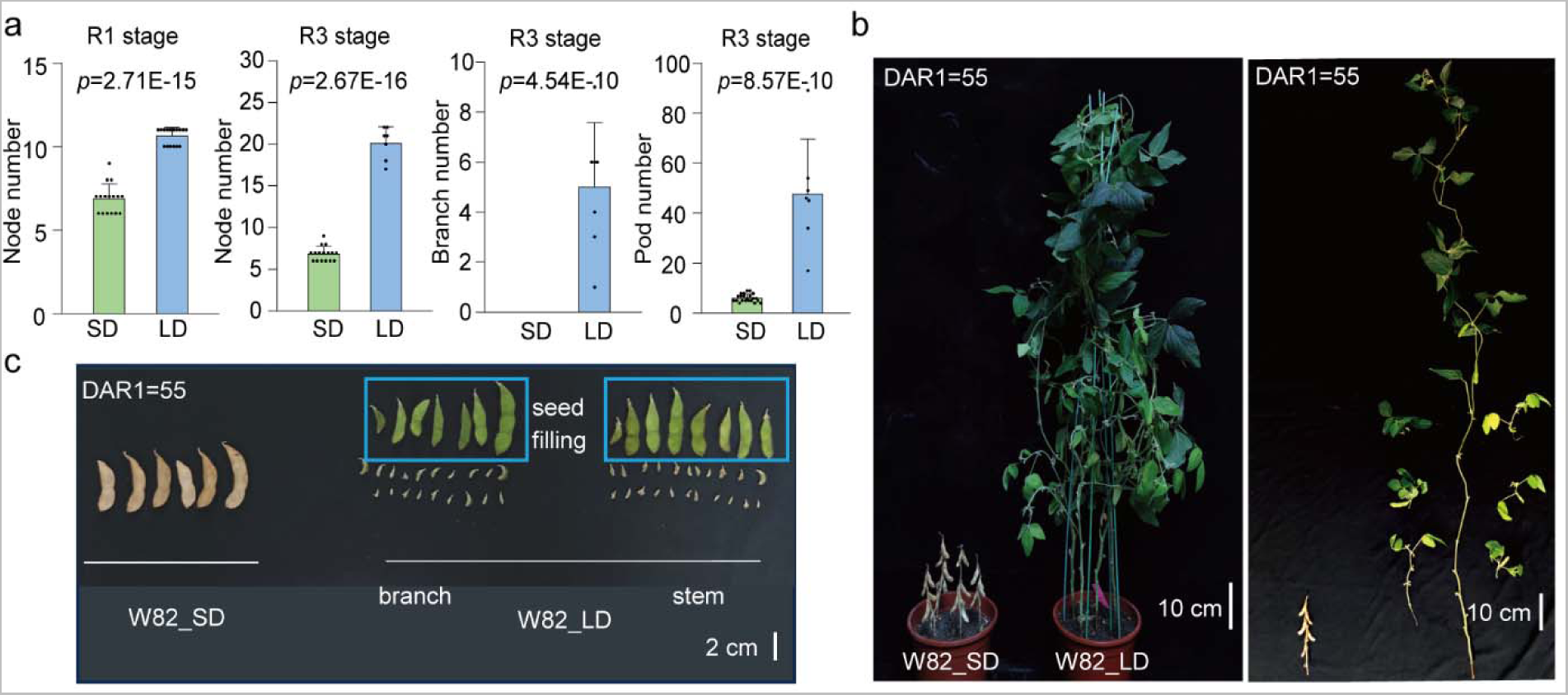
Differences in growth and architecture of soybean cultivar W82 under differing photoperiod conditions. (a) Numbers of nodes on the main stem at flowering stage (R1) and pod setting stage (R3), and pod and branch numbers at R3 of W82 in SD (12 h light/12 h dark) and LD (16 h light/8 h dark) conditions. (b) Phenotype of W82 under SD and LD photoperiod conditions. Seed are sown at the same time, while plants matured faster, were shorter, produced fewer node, and fewer branches under SD than LD. (c) Pod growth status of SD and LD conditions at 55 days after R1 (DAR1). When SD plants reached its maturity stage, total pod number of per plant and developmental stages under two photoperiod conditions were observed at this time-point. Brown pods are ripe, green are unripe. Under SD conditions plants had no branch. And pods on branches and main stem under LD conditions were present here. All data are given as means ± s.e.m.. One-tailed, two-sample t-tests were used to generate the *P* values.

To further observe post-flowering photoperiod sensitivity, we employed a photoperiod transfer experiment, and simulated LD (16 h light/8 h dark) and SD (12 h light/12 h dark) on the two branches of the same decapitated soybean plant. In the photoperiod transfer experiment, the soybean plants of W82 were grown in LD (16 h light/8 h dark) conditions until the R1 stage, after which half were transferred into SD (12 h light/12 h dark) conditions (LD_SD group), with the remaining half continuing to grow under the LD conditions as a control (LD_LD group). Compared to the LD_SD group, the LD_LD group took longer days to initiate podding (Fig. 3). About 14 days after being moved to the SD conditions, the soybean plants of W82 began to successfully set pods but there was no pod setting under continuing LD conditions (Fig. 3a-3b). At 45 days after the photoperiod transfer treatment, pod and seed development under SD conditions were significantly further than under LD conditions, indicating that SD conditions promoted faster development after flowering (Fig. 3c). In the experiment of LD and SD simulation on the same plant, to obtain long branches at similar stages of growth, the SAMs of the soybean plants were removed five days after their emergence under LD conditions (Fig.S3a-b). This released apical dominance, resulting in two symmetrical axillary buds that later developed into two long branches, unlike untreated soybean plants with a single main stem and short branches (Fig. S3c). Next, different photoperiod treatment combinations were applied to the two long branches of each SAM-removed plant after flowering (R1 stage); in the SD&LD combination, one branch was covered with a black plastic bag at ZT12 (LD condition) to simulate the SD condition, with the other branch was covered with a transparent plastic bag to maintain the LD condition, with bags removed daily at ZT0 (Fig. 4a). Under the SD&LD treatment, the branch under the simulated SD conditions set pods earlier than those under the LD conditions, with podding occurring approximately 9 days after shading treatment and reaching the filling stage 15 days after treatment. No pod formation occurred even after prolonged exposure to LD conditions (Fig. 3e, 3i).

**Fig. 3.**
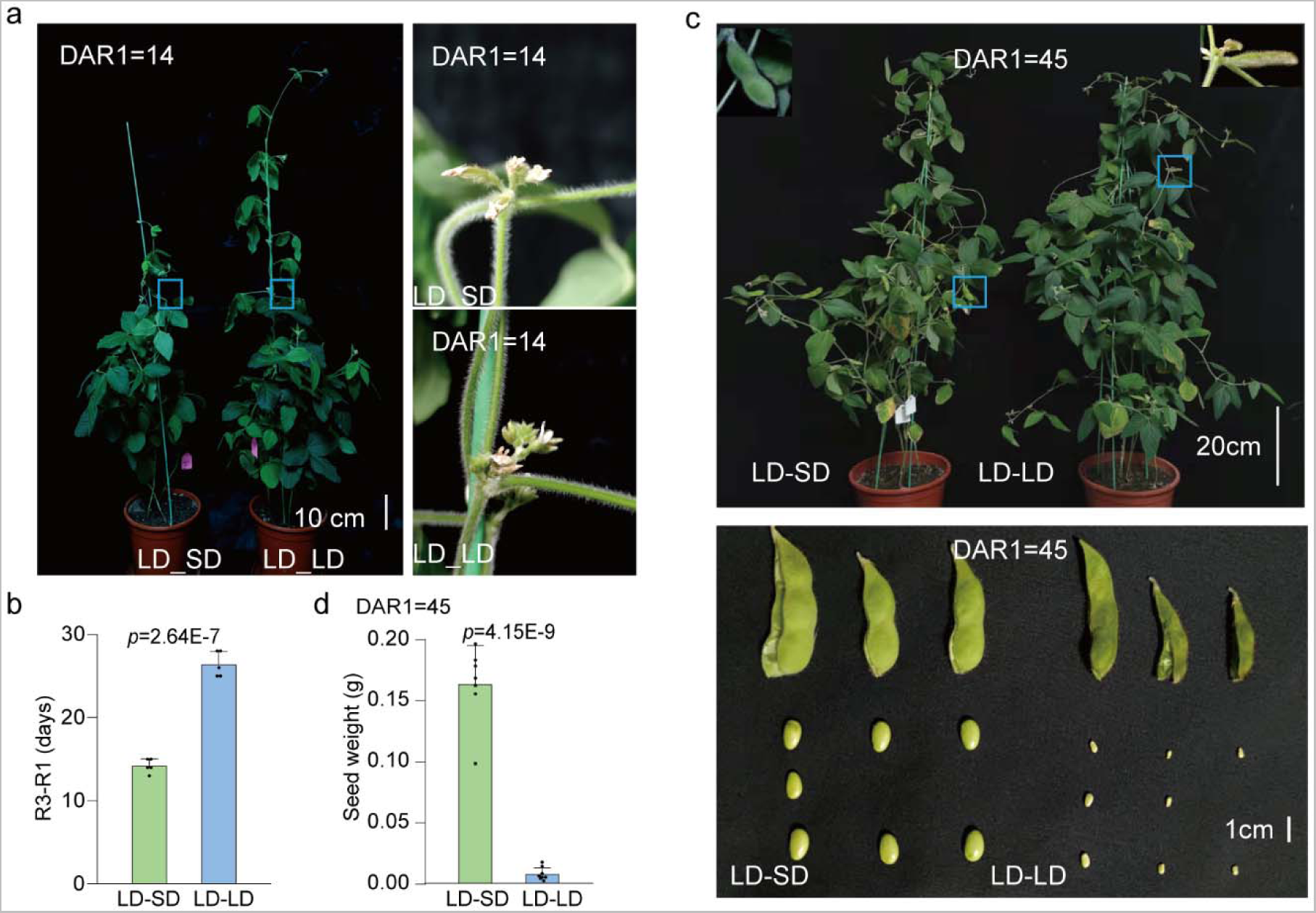
Soybean remains photoperiod sensitive after flowering. In order to detect effect of photoperiod transfer on post-flowering development, half of the 10 plants grown in LD (16 h light/8 h dark) conditions were transferred to SD (12 h light/12 h dark) conditions at the R1 stage (named LD_SD group), while the remaining 5 plants continued to grow under continuous LD condition (named LD_LD; control group). (a) Pod setting was initiated about 14 days after transplantation in the LD_SD plants, but not in the LD_LD conditions. (b) The time required from R1 (time of the first opened flower) to R3 (initiation time of podding) of LD_SD and LD_LD experiment groups. (c) Three representative pod and seed statuses of 45 days after the photoperiod transfer treatment. (d) Fresh seed weight of LD_SD and LD_LD groups at 45 days after the photoperiod transfer treatment. All data are given as means ± s.e.m.. One-tailed, two-sample t-tests were used to generate the *P* values.

**Fig. 4.**
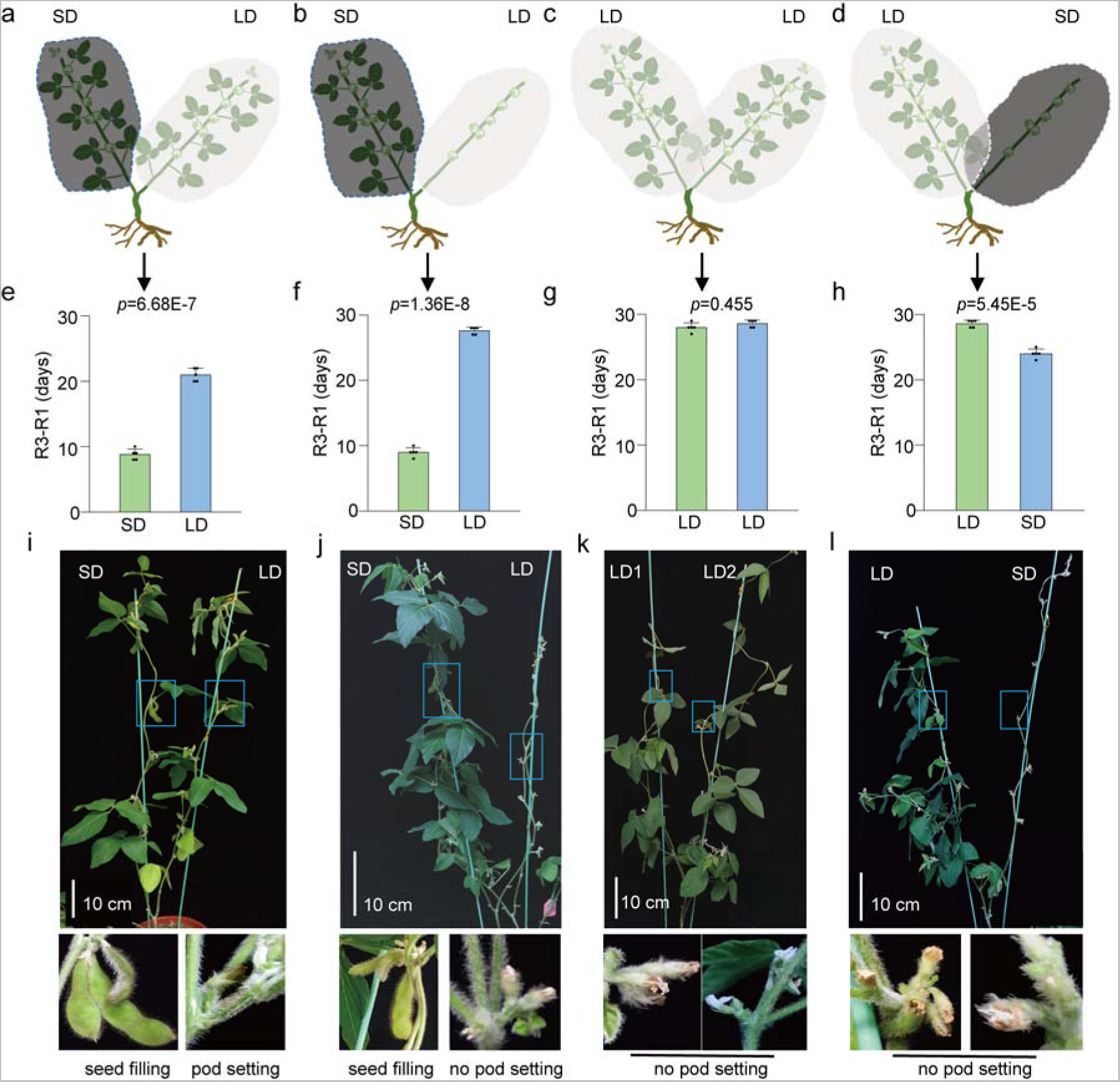
Branch-specific photoperiod treatments reveal that the leaves are responsible for the pod-setting signal. (a) Four groups of branch-specific photoperiod treatments. The shoot apical meristem (SAM) was removed from soybean seedlings in LD (16 h light/8 h dark) conditions, resulting in the simultaneous development of two lateral branches. Different photoperiod treatment combinations were applied to two branches after flowering (R1). (a) SD&LD group: Under normal long-day (LD; 16 h light/8 h dark) conditions, one branch was covered with a black plastic bag at ZT12 to simulate the short-day (SD; 12 h light/12 h dark) condition, while another branch was treated with a transparent plastic bag to maintain the LD conditions. In order to demonstrate that the leaves are the main organs sensing photoperiod and transmitting podding signals, the leaves were also removed from either the SD or LD branches undergoing the SD&LD treatment (b and d). (c) LD&LD group: As a control, both branches were covered with transparent plastic bags to maintain LD conditions. (e–h) Days required from flowering to podding (R3-R1) of four groups in (a-d). (i-l) The phenotypes of the different photoperiod combinations described in (a-d) at 15 days after treatment. Branches in the SD condition with leaves successfully set pods, and at 15 days the pods had reached the seed-filling stage (i and j). Under LD the pod-setting time was later than that of the SD, whether or not the leaves were removed. All data are given as means ± s.e.m.. One-tailed, two-sample t-tests were used to generate the *P* values.

These results show that soybean remained sensitive to photoperiod even after flowering, especially reflected in different pod-setting times, suggesting that plant architecture may not be the sole factor contributing to this difference.

### The photoperiod-regulated pod-setting signal is mainly induced in and transmitted within the leaves

How photoperiod affects the conversion of open flowers to pods or shedding? We set different photoperiodic conditions for branches on the same plant, in addition, to prove that leaves are the main organs for perceiving photoperiod and transmitting podding signals, we removed the leaves of the branches under the different photoperiod conditions of the SD (12 h light/12 h dark) &LD (16 h light/8 h dark) treatment (Fig. 4a-4d). For this study, 5-day-old soybean seedlings were decapitated at cotyledon stage, and there were no leaves from other parts of soybean except the two branches. Our treatment included SD&LD, SD&LD (with no leaves after R1 under LD condition), LD&LD and LD&SD (with no leaves after R1 under SD condition) four experimental groups. The LD&LD combination was a control, in which both branches were covered with transparent plastic bags (Fig. 4c). The pod-setting time under the SD conditions was prolonged by removing the leaves in LD&SD (with no leaves after R1) group (Fig. 4e, 4h). Under LD conditions, the pod-setting time was longer than under SD conditions, regardless of leaf removal (Fig. 4e–4h). However, comparing the branches at stage R3 under LD conditions in the four groups, we found that the onset of pod formation in the LD&SD treatment group occurred approximately one week earlier (about 21 days) than in the other three groups (about 30 days) (Fig.4e-4h). The pod formation signal should be perceived by the leaves, transmitted downward, and communicated between different branches. Moreover, the signal inducing short-day pod formation is stronger than that promoting long-day flower abscission. From these results, we infer that leaves are the main light sensor and that the photoperiod signal is mainly induced in the leaves, which then transmit the signal to form pods to the flowers.

### *E1* is downstream of the EC in controlling pod-setting time

As reported that the homologs of *PHYA*, members of the evening complex (*EC*), *E2* and *E1* are the major genetic players in the control soybean photoperiod sensitivity, and their functions are mainly described in regulating flowering time (Lin *et al*., 2022; Bu *et al*., 2021; Zhao *et al*., 2024). To further explore the genetic pathway underlying how the photoperiod affects the pod-setting time after flowering, we investigated the pod-setting time of photoperiod-insensitive mutants under different photoperiod conditions. The early-flowering triple mutant *e1-3m*, which is insensitive to photoperiod, underwent early pod setting after flowering, with no differences under different photoperiods (Fig. S4a-4c). The late-flowering double mutant *lux-2m*, which was also insensitive to photoperiod, had later flowering times and pod-setting times than the wild type under particularly long day (20 h light/4 h dark) conditions (Fig. S4d, 4e). To examine whether the difference in flowering and podding times between the wild type and late-flowering mutants disappears under extremely long photoperiods, we selected exceptionally long photoperiods. *e1-3m* was crossed with *lux-2m* to obtain *e1-3m lux-2m* quintuple mutant. Under LD (16 h light/8 h dark) conditions, *e1-3m lux-2m* the multiple mutants showed early pod setting, which was similar to the *e1-3m* phenotype (Fig. S4f). *Luxs* are parts of the *EC* in circadian clock (Nusinow *et al*., 2011; Jung *et al*., 2020; Bu *et al*., 2021). This indicates that *E1* is downstream of the *EC* in controlling the initiation of pod setting, and that pre- and post-flowering photoperiodic sensitivity may be controlled by the same genes. However, after the input of photoperiodic signal, the response genes controlled different development processions may be different.

### Photoperiod affects pistil development

In previous experiments, we found that flowers opened in LD conditions tend to falling before pod formation. We sought to investigate whether there are differences in pollen viability and pistil morphology between LD and SD conditions, leading to differences in pod formation time. We sought to investigate whether there are differences in pollen viability and pistil morphology between LD and SD conditions, leading to differences in pod formation time. We collected pollen from W82 under LD and SD conditions and conducted pollen germination experiments *in vitro*. Results revealed no significant differences in pollen tube length and pollen germination rate (Fig. S5a-5d). Additionally, unopened flower buds under LD and SD conditions were emasculated and artificially pollinated, and pollen tubes were able to germinate normally *in vivo* (Fig.S5e-5f). The effect of photoperiod on pollen viability may be minimal. We found that there were morphological differences in pistil morphology under LD and SD conditions (Fig. 5a). This morphological difference leads to the similar height of pistil and stamen when stamen begins to disperse powder under short-day conditions (Fig. 5b-5c, Fig. S6a, 6c-6d), facilitating rapid and successful pollination. While the height of stamen is lower than that of pistil under long-day conditions (Fig. 5b-5c, Fig. S6a, 6c-6d), which is not conducive to rapid pollination. Flower buds or open flowers exhibit similar external sizes and shapes under both LD and SD conditions, but significant differences exist in stigma sizes. (Fig. S6a-6b, Fig. S6e-6g). The morphology of pistil styles varied greatly in the late development stage of buds. Under SD conditions, a hook-like structure is present at the apex of the stigma, whereas under LD conditions, the curved hook is less pronounced. And when moved from LD to SD for a period of time, the hook structure at the apex of newly emerged flower buds becomes pronounced (Fig. 5a-5c, Fig. S7).

**Fig. 5.**
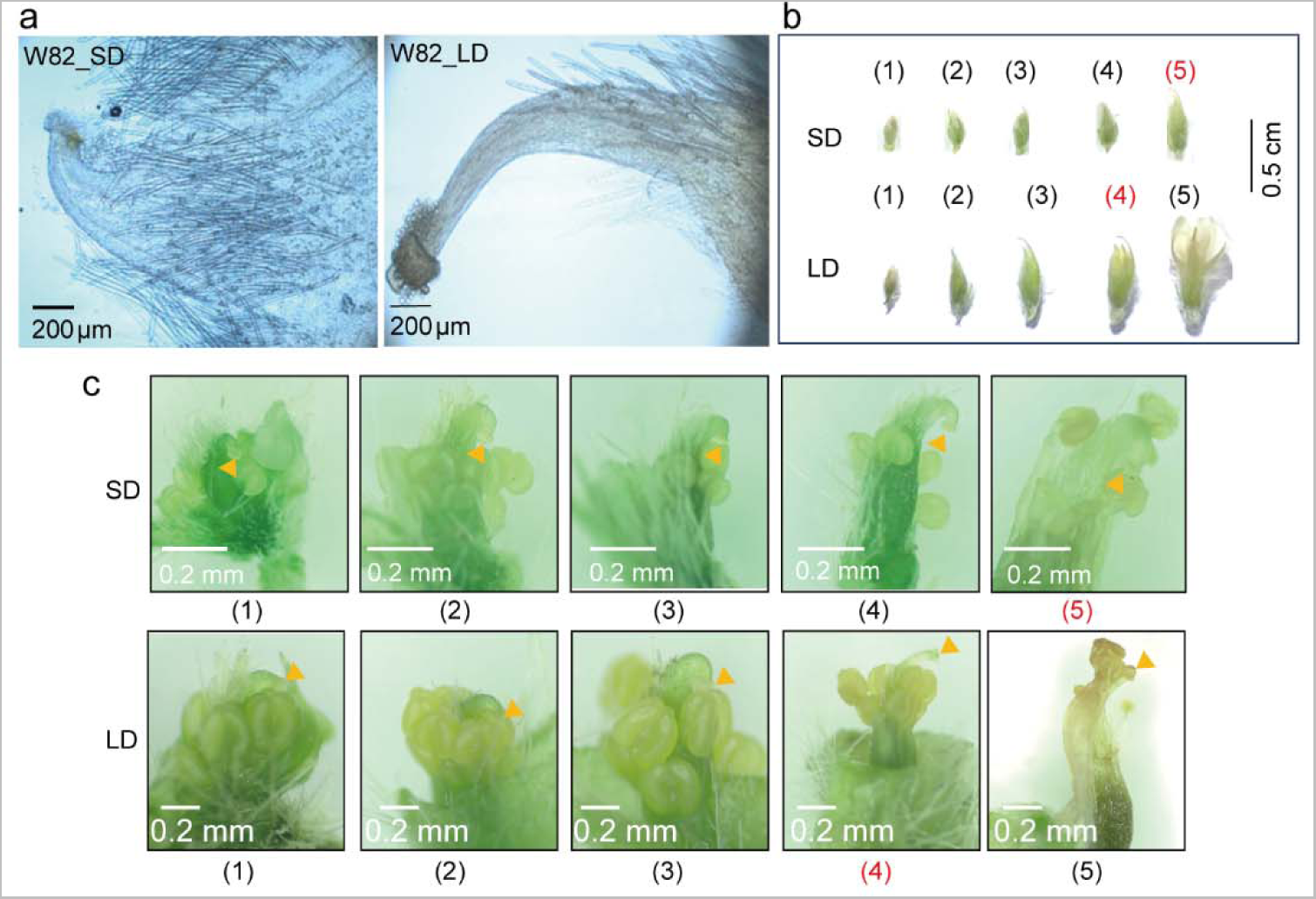
Photoperiod affects style morphology and the development of pistil and stamen of soybean. (a) Phenotype of the styles of opened flowers of W82 under SD (12 h light/12 h dark) and LD (16 h light/8 h dark) conditions, respectively. (b) Flowers or buds in different development stages in an inflorescence under SD and LD conditions of W82. (c) Growth status of pistil and stamen of bud or flower in (b). The numbers marked in red represent the buds with pollen grains dispersed from anthers. The orange triangle represents the position of the stigma.

Which genes and plant hormones effect pistil development under different photoperiod conditions? We collected flower buds under LD and SD conditions, measured plant hormone levels, and isolated pistils for RNA extraction, constructing RNA-Seq libraries and analyzing differentially expressed genes. Simultaneously, we analyzed the relative expression levels of differentially expressed genes in the buds of the top three nodes of soybean plants under the continuous long day (LD_LD) and LD_SD groups at R1, 1 day, 5 days, and 10 days after the R1 stage. According to the sequencing results (Supplementary table 1), the regulatory pathways of differentially expressed genes involve the MAPK signaling pathway, starch and sucrose metabolism, photosynthesis, plant hormone signal transduction (Fig. S8). We identified at least 23 DEGs might affect soybean pod formation (Fig. 6a, Supplementary table 2). We selected 9 genes from the 23 DEGs for PCR verification. Consistent with our qRT-PCR analysis (Fig. S9), *REPRESSOR OF PHOTOSYNTHETIC GENES 2* (*RPGE2*), *GIBBERELLIN* OXIDASE *8* (*GA2OX8*), and *GA2OX2* were upregulated upon transfer to SD conditions, while *WRKY19*, *RESPIRATORY BURST OXIDASE HOMOLOGUE E* (*RBOHE*), *RBOHB*, *SUCROSE PHOSPHATE SYNTHASE 3F* (*SPS3F*), and *Xyloglucan Endotransglucosylase/hydrolase*s *(XTHs*) were strongly inhibited (Fig. 6a, Fig. S9). As expected, the content of some plant hormones varied in the buds under LD and SD conditions (Fig. 6b, Supplementary table 3). The contents of gibberellin 1, 3, 7 (GA1, GA3, and GA7), cytokinin and salicylic acid were higher under LD condition. The contents of auxin and jasmonic acid were higher under SD condition. Under LD conditions, after flowering (R1 stage), a 50 mg/ml sucrose solution was applied on the leaves, and the control group was sprayed with the same amount of water. The results showed that the external application of sucrose solution could promote pod formation (Fig. 6c). All these results suggest that photoperiod may control soybean pod formation and development by regulating multiple gene pathways and plant hormones.

**Fig. 6.**
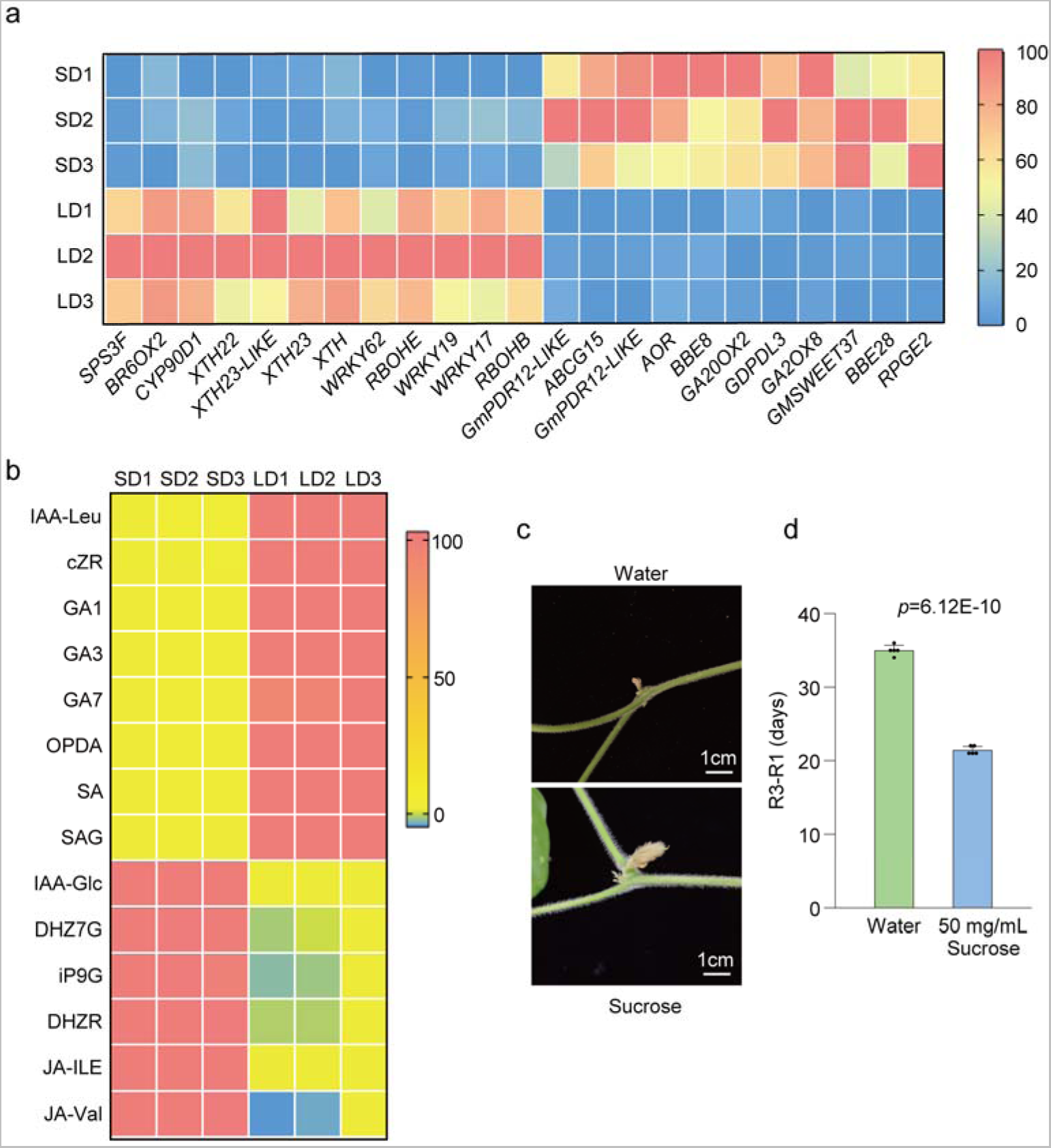
Comparison of transcripts activities and plant hormone contents in pistils of SD (12 h light/12 h dark) and LD (16 h light/8 h dark) conditions. (a) Some differentially expressed genes in pistils of SD and LD conditions. (b) Some plant hormones with significant differences in content of SD and LD conditions. (c) Pod or flower morphology at the fourth upmost node of control groups and the external application of 50 mg/mL sucrose solutions groups. (d) External application of sucrose shortens the time required for initial pod setting under LD conditions. All data are given as means ± s.e.m.. One-tailed, two-sample t-tests were used to generate the *P* values.

## Discussion

Photoperiod regulates various growth and development processes of such as floral induction and stem termination and pod development in post-flowering reproductive growth stage (Han *et al*., 2006; Xu *et al*., 2013; Nico *et al*., 2016). Previous studies have found that exposing soybean plants to long-day conditions during post-flowering reproductive growth stages extends the R3-R6 period, with seed development and seed number positively correlated with the duration of the R3-R6 stage (Kantolic *et al*., 2001; Kantolic *et al*., 2007). These results indicate that soybean plants remain sensitive to photoperiod during the post-flowering R3-R6 stages. In this study, we found that different soybean cultivars are sensitive to photoperiod in initiation of podding after flowering in laboratory-controlled conditions. SD conditions promote pod formation while LD prolonged duration of R1 to R3 stage. The photoperiod-insensitive mutants used in this study might provide a basis for further research on the mechanism of photoperiod-sensitive related genes in regulating the pod-initiation time. The photoperiod-insensitive *lux-dm* and *e1-3m* mutants (Bu *et al*., 2021; Lin *et al*., 2022) display two extreme phenotypes. The *lux-dm* mutants had late flowering time, produce more stem nodes, branches, and leaves than wild-type soybean plants (Bu *et al*., 2021; Lin *et al*., 2022), while the *e1-3m* mutant had a smaller morphology with few nodes and early flowering time (Lin *et al*., 2022). In this study, we found that *e1-3m* had short R1-R3 stage (about 5 days) in both LD and SD conditions, pod initiation time was non-sensitive to photoperiod. The *lux-dm* mutant exhibited a longer R1-R3 duration. While *e1 e1la e1lb lux1 lux2* quintuple mutant showed an R1-R3 duration similar to e1-3m, suggesting that E1 and E1-likes function downstream of the EC in controlling pod-setting time, with EC being entirely dependent on E1. The mechanisms of photoperiod signal sensing and transmission may remain conserved before and after flowering. *E1*, *E1-likes*, and *EC* have been reported to play major roles in floral induction (Lin *et al*., 2022, Bu *et al*., 2021, Xia *et al*., 2012), but their roles in post flowering reproductive development remain undetermined. Increasing research attention is being given to the effects of growth period genes on post-flowering development (Takeshima *et al*., 2019, Wan *et al*., 2022).

The coordination of flower development and fertility is regulated by endogenous developmental signals such as the phytohormones jasmonates (JAs), auxin, and gibberellin, as well as environmental cues. (Huang *et al*., 2023). We found that under LD photoperiod conditions, the firstly-opened flowers typically dropped, and the second-round flowers slowly turn into pods. In our study, we found that pistil style of W82 exhibit different morphology, when the anther of the stamen was dispersed, the stigma is higher than that of the stamen. Under SD conditions there were apical hook formations in flower style like hook in emerging seedlings. Longer style length in rice influences the stigma exertion and increase outcross rate of male sterile line and the yield of hybrid F1 seeds. The elongation of cell length in the style is associated with a higher GA4 content in the pistil (Dang *et al*., 2022). We found that under LD conditions endogenous GA1, GA3, and GA7 content in flower buds were higher than that in SD conditions, but lower IAA-Glc and JA-Ile content. Apical hook formation involves a gravity-induced auxin maximum on the eventual concave side of the hook (Du *et al*., 2022). Jasmonoyl-L-isoleucine (JA-Ile) is a biologically active form of JA. JA-deficient mutants exhibit low fertilization rates and abnormal flower formation (Riemann *et al*. 2008, 2013, Cai *et al*. 2014, Xiao *et al*. 2014, Hibara *et al*. 2016, Inagaki *et al*., 2023). The *jasmonic acid insensitive 1-1* (*jai1-1*) mutants in tomato exhibits arrested flower bud development just before flower opening by abolishing the peaks of JA biosynthesis and *SlMYB21* expression in flower buds within ∼2 d before flower opening (Dobritzsch *et al*., 2015; Niwa *et al*., 2018). These results suggest that JA plays a crucial role in flower development and fertility in rice and tomato. We performed RNA-seq on pistils of W82 flower buds under LD and SD. Compared to pistils under LD condition, plant cell wall remodeling enzymes *XTH22*, *XTH23*, and *XTH23-like* genes were significantly decreased in SD conditions (Fig.6a, Fig.S9), XTH22 and XTH23 are known to play a role in cell elongation during flower development (Claisse *et al*., 2007). *RbohB* and *RbohE* genes were up-regulated in LD conditions. Upon transition from LD to SD, their relative expression levels were down regulated (Fig.6a, Fig.S9). RBOHs are reported to be crucial for ROS generation and are essential for precise flower and fruit abscission (Lee *et al*., 2018, Ma *et al*., 2023). Previous studies have shown that under a photoperiod of approximately 14.5 hours of light per day, about 21%-28% of flowers and pods are aborted, which increases to 42%-49% with shading treatments (Ali *et al*., 2022). Top bud removal at each node, leaving only one remaining top bud, can reduce flower abscission rates, while shading treatments do not increase flower abscission rates. Bud removal at each node, leaving only one remaining bud, can reduce flower abscission rates, while shading treatments do not increase flower abscission rates (Ali *et al*., 2022). This suggests that light/shade conditions are not directly responsible for flower/pod abscission signals; rather, a lack of nutrient supply leads to increased flower and pod abscission rates (Ali *et al*., 2022). In our study, we found that even under SD condition that promoted pod setting, pod formation could not be achieved as rapidly after leaf removal as it was in the experimental group that retained its leaves (Fig. 3d, 3h, 3I), likely because photosynthesis and assimilate accumulations were decreased. Enhanced carbon assimilation could reduce flower and pod abortion, as well as accelerating leaf expansion, seed yield, and the production of tuberous storage organs or fibers in various crops (Abelenda *et al*., 2019; Ali *et al*., 2022; Yue *et al*., 2021; Xu *et al*., 2012). In this study, KEGG enrichment analysis of the DEGs in the buds before opening of soybean revealed that genes related to starch and sucrose metabolism, carbohydrate or energy metabolism were repressed under LD conditions. Application of exogenous sucrose solution promoted pod formation. It has been reported that during the early stages of seed development, embryos grow rapidly and acquire a large amount of sugar from liquid endosperm. An insufficient supply of nutrients from the endosperm to the embryo results in severe seed abortion and yield reduction (Wang *et al*., 2019). Soybean seed development responds to photoperiod, where the Dt1 protein physically interacts with the sucrose transporter GmSWEET10a, negatively regulating the transport of sucrose from seed coat to embryo, thus modulating seed weight under LD conditions. *Dt1* exhibits pleiotropy in regulating both seed size and stem growth habit in soybeans (Li *et al*., 2024). The photoperiod-insensitive mutants used in the present study might provide a basis for further studies into the mechanism by which the photoperiod-sensitive flowering pathway genes regulate the pod-initiation time and pod number through the photoperiod-dependent regulation of the balance between source and sink tissues.

## Supplementary Data

Fig. S1 The difference in pod initiation time after flowering of Williams 82 (W82) under long-day (LD) and short-day (SD) conditions in greenhouse.

Fig. S2 Under different photoperiod conditions, the growth and development process of soybean.

Fig. S3 Decapitation treatment of soybean plant.

Fig. S4 The pod initiation time of the triple mutant (*e1/e1la/e1lb*, *e1-3m*) is insensitive to photoperiod.

Fig. S5 Effect of photoperiod on pollen germination. Fig. S6 Photoperiod affect pistil and stamen growth.

Fig. S7 Photoperiod influence the morphology of flower style.

Fig. S8 Enrichment analysis of differentially expressed genes in pistil of W82 under LD (16 h light/8 h dark) and SD (12 h light/12 h dark) conditions.

Fig. S9 Relative expressions of pistil growth and development related genes in W82 under LD_LD and LD_SD conditions at different time.

Fig. S10 A proposed working model for SD (12 h light/12 h dark) and LD (16 h light/8 h dark) regulate soybean photoperiod podding.

Supplementary table 1 A total of 5239 DEGs in pistils of Williams 82 between long-day and short-day conditions.

Supplementary table 2 The 23 genes of the 5239 DEGs of Williams 82 between long-day and short-day conditions.

Supplementary table 3 Phytohormones contents in flower buds under long-day and short-day conditions.

Supplementary table 4 Primers for quantitative RT-PCR.

## Acknowledgments and funding

This work was supported by the National Natural Science Foundation of China (grant 31901569 and 32372193 to L.K.).

## Author contributions

F.K., B.L., and S.L. supervised the experiments. L.K., Y.W., H.L. and X.H. performed the research. L.K. analyzed the data with the help of R.F., L.Y., and Y.W.. L.K. and Z.S. wrote the draft manuscript with the input from X.L.and S.L.. H.L. and L.C. assisted in editing the manuscript. All authors read and approved the final manuscript.

## Data availability

Data supporting the findings of this study are available in the supplementary material of this article.

